# Rapid and Stimulus-Specific Deviance Detection in the Human Inferior Colliculus

**DOI:** 10.1101/2024.06.18.599524

**Authors:** Johannes Wetekam, Nell Gotta, Luciana López-Jury, Julio Hechavarría, Manfred Kössl

**Author notes:** Corresponding authors: Johannes Wetekam, Julio Hechavarría. Shared last authorship (Julio Hechavarría, Manfred Kössl).

## Abstract

Auditory deviance detection, the neural process by which unexpected stimuli are identified within repetitive acoustic environments, is crucial for survival. While this phenomenon has been extensively studied in the cortex, recent evidence indicates that it also occurs in subcortical regions, including the inferior colliculus (IC). However, compared to animal studies, research on subcortical deviance detection in humans is often constrained by methodological limitations, leaving several important questions unanswered. This study aims to overcome some of these limitations by employing auditory brainstem responses (ABRs) to investigate the earliest neural correlates of deviance detection in humans, with a focus on the IC. We presented healthy participants of either sex with low- and high-frequency chirps in an oddball paradigm and observed significant deviance detection effects in the ABR, specifically when low-frequency chirps were used as deviants within a context of high-frequency standards. These effects manifested as larger and faster ABRs to deviant stimuli, with the strongest responses occurring at higher stimulation rates. Our findings suggest that the human IC exhibits rapid, stimulus-specific deviance detection with differential modulation of response amplitude and latency. The data indicate that the temporal dynamics of novelty detection in humans align well with the data reported in animals, helping to bridge the gap between animal and human research. By uncovering previously unknown characteristics of subcortical deviance detection in humans, this study highlights the value of ABR recordings with excellent temporal resolution in investigating subcortical deviance detection processes.

**Significance statement:** Auditory deviance detection enables the brain to identify unexpected stimuli in a repetitive environment, but its subcortical mechanisms in humans remain comparatively underexplored. Using auditory brainstem responses (ABRs), our study reveals two key findings about deviance detection in the human inferior colliculus (IC). First, we show subcortical deviance detection at latencies under 10 ms, bridging a longstanding gap between human and animal research. Second, deviance detection in the IC is rapid, emerging within three or fewer standard repetitions, with differential modulation of ABR amplitude and latency. These findings improve our understanding of the temporal dynamics of auditory processing in the human IC and highlight the value of ABR recordings in studying subcortical deviance detection mechanisms.

## 1. Introduction

The ability of the brain to detect unexpected acoustic stimuli within a repetitive environment, known as auditory deviance detection, has been extensively studied. This phenomenon has been characterised in detail through animal and human research (Näätänen et al., 2007; May and Tiitinen, 2010; Escera and Malmierca, 2014; Carbajal and Malmierca, 2018). Initially thought to be limited to the cortex (Näätänen et al., 1978; Ulanovsky et al., 2003, 2004), studies in rodents revealed that deviance detection is widespread throughout the brain, including several subcortical regions (Kraus et al., 1994a, 1994b; Pérez-González et al., 2005; Malmierca et al., 2009; Antunes et al., 2010; Zhao et al., 2011). Subsequent research demonstrated that correlates of deviance detection are also present in the subcortical auditory system of humans (Chandrasekaran et al., 2009; Skoe and Kraus, 2010; Slabu et al., 2012; Cacciaglia et al., 2015), suggesting a widespread novelty detection mechanism in the mammalian brain (Escera et al., 2014; Parras et al., 2017).

Despite these advances, evidence for subcortical deviance detection in humans is confined to studies using the frequency-following response (FFR) or functional magnetic resonance imaging (fMRI), both of which have poor temporal resolution. This limitation has hindered a detailed understanding of the timescales at which deviance detection operates in the human brainstem. By contrast, animal studies have shown that subcortical stimulus-specific adaptation (SSA), a single-neuron correlate of deviance detection, emerges rapidly within fewer than 10 standard repetitions and that deviant stimuli elicit more frequent and earlier spiking activity compared to standards (Malmierca et al., 2009; Zhao et al., 2011).

In human research, a promising non-invasive technique for overcoming temporal limitations is the recording of auditory brainstem responses (ABRs). ABRs, population-level responses from the brainstem, offer excellent temporal resolution and have been successfully used to study subcortical deviance detection in animals (Duque et al., 2018; Wetekam et al., 2022, 2024). However, attempts to demonstrate deviance detection in human ABRs have so far been unsuccessful (Slabu et al., 2010; Althen et al., 2011), possibly because previous studies examined multiple components of the auditory evoked potential simultaneously, preventing the optimisation of parameters for ABRs.

The current study addresses this issue by optimising stimulation parameters based on insights from previous work. The primary goal is to quantify the dynamics of human subcortical deviance detection, with a particular focus on the timescales at which it operates in the auditory brainstem. ABRs were recorded using low- and high-frequency chirp stimuli (Fig. 1a), designed to maximise ABR amplitude (as introduced by Fobel and Dau, 2004), while maintaining distinct frequency differences between deviant and standard stimuli in an oddball paradigm (Fig. 1b). Animal studies have shown that the effect size of deviance detection increases with higher stimulus repetition rates (Zhao et al., 2011; Patel et al., 2012; Wetekam et al., 2022). To incorporate this finding, ABRs in this study were recorded at three repetition rates, all significantly faster than those used in previous research on ABR deviance detection in humans. The high temporal resolution of ABRs allowed for the analysis of changes in response amplitude and peak latency, as well as the investigation of how subcortical deviance detection modulates sequences of standard responses following a deviant. This analysis provides insights into the number of stimulus repetitions required for deviance detection to emerge in the brainstem.

**Figure 1:**
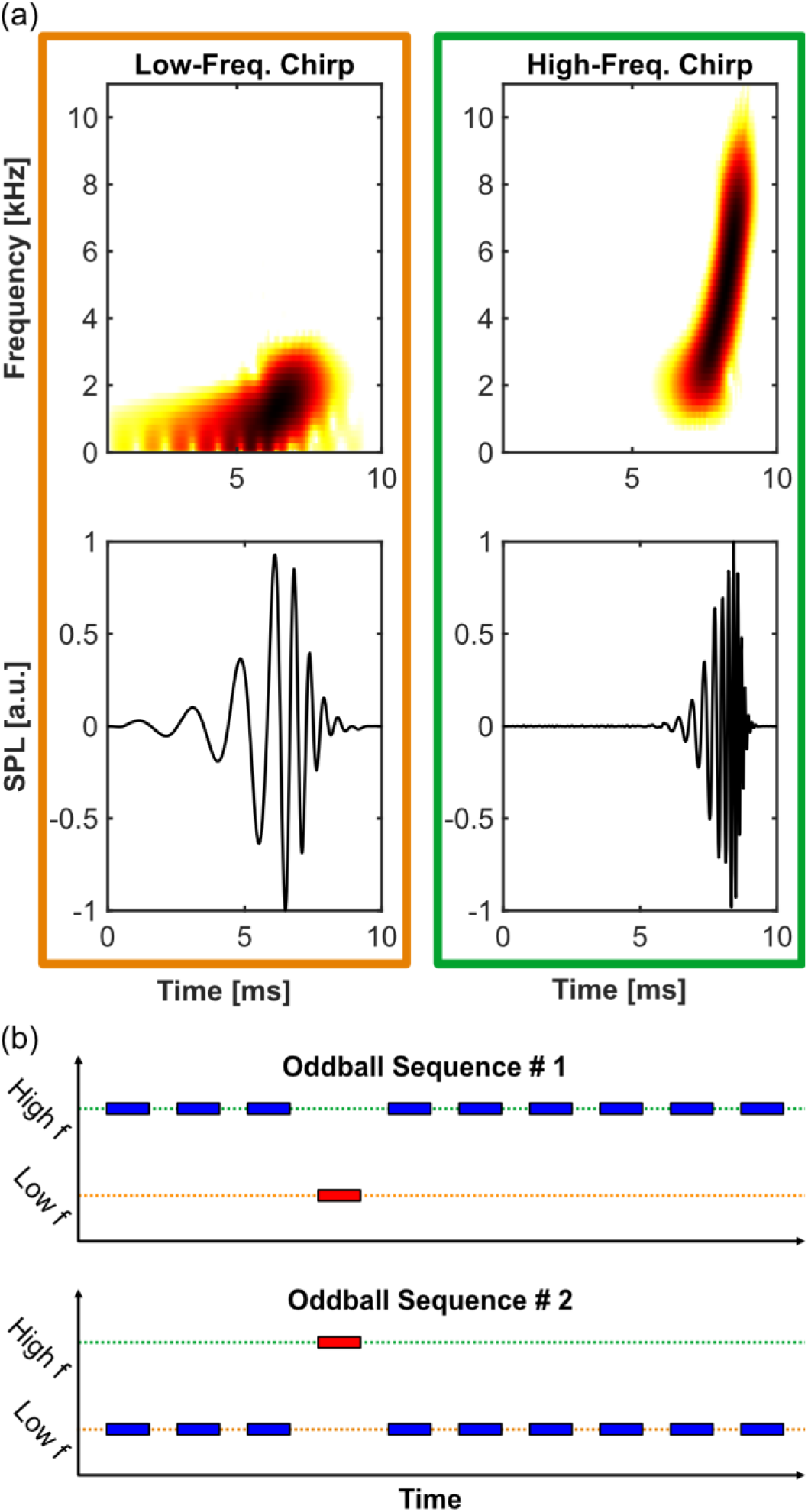
Used stimuli and stimulation protocol. (a) Spectrograms (top) and oscillograms (bottom) of a low-frequency chirp (orange frame) and a high-frequency chirp (green frame) that were used as stimuli in this study. (b) Schematic representation of the oddball paradigm; blue: high-probability standard, red: low-probability deviant.

Based on findings from animal research, we hypothesised the following:

1. Optimising stimulation parameters will enable the measurement of deviance detection in human ABRs.
2. Deviant stimuli will elicit ABRs that are larger and earlier than those to standard stimuli.
3. The modulatory influence of deviance detection will depend on the stimulus repetition rate, with stronger effects at faster rates.
4. Deviance detection will emerge rapidly, within just a few standard repetitions.

The results of this study confirmed all four hypotheses, offering new insights into the mechanisms underlying human subcortical deviance detection. The findings also help bridge the gap between human and animal research in this domain.

## 2. Material and Methods

### Participants

ABRs of 27 participants (8 males, 19 females), aged between 20 and 40 years (mean age = 25.5 years) and with normal hearing, were recorded for the current study. Informed consent was obtained from each participant prior to recording, and all subjects received monetary compensation for their participation. The study was approved by the Goethe University Frankfurt Ethics Committee (reference number: E 27/23) and was conducted in accordance with the WMA Declaration of Helsinki. Before the recording began, a two-alternative forced choice test was used to determine each participant’s individual hearing thresholds for the stimuli used during the ABR recording. Each recording session lasted between 3 and 3.5 hours. Due to limitations of total measurement duration, the data presented in this study was collected in two separate recording sessions. While some participants attended both sessions, others participated in only one. Consequently, the total number of participants exceeds the number of datasets available for analysis.

### Stimulation and recording procedure

Two artificially generated chirps, differing in their frequency composition, were used as stimuli (Fig. 1a). These chirps were designed to maximise ABR amplitudes by compensating for the cochlear traveling wave delay, as described by Fobel and Dau (2004), Elberling et al. (2007) and Elberling and Don (2008). The chirp design was based on the latency-frequency function derived from narrowband ABR data by Don et al. (2005). These chirp stimuli effectively evoked a prominent IC-generated wave V (Buchwald and Huang, 1975; Henry, 1979; Caird and Klinke, 1987; Land et al., 2016), thereby maximising the overall ABR amplitude. However, the earlier ABR components (waves I-IV) were typically less pronounced compared to those evoked by click stimuli. The stimuli were created with a logarithmic frequency distribution: the low-frequency chirp ranged from 500 Hz to 2 kHz, and the high-frequency chirp ranged from 2 kHz to 8 kHz. Importantly, although both chirps had different durations (low-frequency chirp: 9 ms, high-frequency chirp: 4 ms), their design ensured a simultaneous activation of the entire basilar membrane, occurring temporally close to the peak of each stimulus. This characteristic, combined with the fact that ABRs reflect onset responses and comparisons were made exclusively between responses to physically identical stimuli, makes it highly unlikely that the difference in stimulus duration has influenced the results of this study. The signals were digitally generated using custom-written Matlab (MathWorks) scripts, converted from digital to analogue by a 192 kHz Fireface UC soundcard (RME Audio), and delivered to the participant’s right ear via an ER-3C earphone (10 Ω, Etymotic Research).

During the ABR recording, stimuli were presented to the participants at 40 dB HL, in two pseudo-randomly arranged oddball sequences. In the first sequence, the high-frequency chirp was presented as high-probability standard (90 %, 9000 repetitions), while the low-frequency chirp was presented as low-probability deviant (10 %, 1000 repetitions). The sequence was then repeated with the roles of the deviant and standard stimuli swapped (Fig. 1b), resulting in two distinct responses for each stimulus: one when perceived as the standard and one as the deviant. The recordings were conducted at three different stimulus presentation rates: 40 Hz, 33 Hz and 20 Hz. Due to constraints on total measurement duration, each recording session included either the two faster rates (40 Hz and 33 Hz) or the slower rate (20 Hz). In sessions involving the two faster rates, the order of their presentation was randomised across participants.

ABRs were recorded using two Ag-AgCl electrodes placed on the participant’s skin: one at the vertex (Cz, active) and the other on the ipsilateral earlobe (reference). A third electrode on the forehead served as the ground. Impedance was reduced to less than 10 kΩ, and data was collected at a sampling rate of 10 kHz. The recordings were conducted in an electrically and sound-insulated booth, where participants sat comfortably in a reclining chair. They were instructed to relax, keep their eyes closed to prevent eye-blink artefacts, and minimise movement during the recording.

### Data processing and statistical evaluation

All data analysis and statistical evaluation were conducted using Matlab. The recorded data was bandpass filtered using a 70-1500 Hz Butterworth filter (4th order). This relatively narrow filter was specifically chosen to effectively remove slow-frequency middle-latency response (MLR) components from the signal, which could otherwise overlap with the ABR of the subsequent stimulus. Such overlap would have confounded the analysis, as differences between deviant and standard ABR amplitudes may then not truly reflect sensitivity to stimulus probability but could instead arise from differences between the preceding low- and high-frequency MLR components. For baseline correction, the mean response amplitude of each trial was calculated within a 5 ms time window immediately before ABR onset. Since the chirp stimuli used in this study were designed to activate all regions of the basilar membrane simultaneously, lower frequencies were presented before higher frequencies in each chirp, compensating for the basilar membrane’s travelling wave delay (Fobel and Dau, 2004; Elberling et al., 2007). As a result, the chirp onset does not coincide with the onset of basilar membrane stimulation, as it would for pure tone stimuli. Therefore, rather than using the stimulus onset as the end of the baseline correction window, it was aligned with the time point when the maximum stimulus amplitude reached the participant’s ear, corresponding to the actual onset of basilar membrane stimulation. The mean amplitude value within this window was then subtracted from the entire trial, resulting in a mean pre-ABR activity of 0 µV.

The first part of analysis in this study aimed to determine whether the recorded ABRs were generally sensitive to the stimulus’ probability of occurrence. To achieve this, the averaging process included only standard trials immediately followed by a deviant trial, and only deviant trials immediately preceded by a standard. This approach ensured that the same number of deviant and standard trials were used to calculate the average response for each participant (ranging between 851 and 914 trials) in order to maintain equal variance across conditions. Moreover, by focusing on standard responses at the end of a sequence (just before a deviant), this analysis maximised the potential to detect differences between deviant and standard responses. All trials were corrected for the sound travel delay (0.76 ms) caused by the distance between the speaker and the ear (26 cm). In each time-voltage graph, t = 0 represents the time point when the maximal stimulus amplitude reached the ear.

To identify differences between deviant and standard ABRs, a cluster-based permutation analysis was performed (in accordance to Candia-Rivera and Valenza, 2022). First, a Wilcoxon rank-sum test was used to compare the deviant and standard ABRs at each time point. If more than five consecutive time points had p-values below 0.01, these points were grouped into a cluster. Next, the z-values for each time point within a cluster were summed to obtain the cluster weight. To determine whether the observed clusters represented true effects of deviance detection or were simply due to random variations in the data, a permutation distribution was generated. Deviant and standard trials were randomly shuffled, and the average ABR waveforms were recalculated, producing two new shuffled responses for each subject. Clusters and their weights were then recomputed. This permutation procedure was repeated 1000 times, creating a distribution of cluster weights under the null hypothesis that no real difference exists between deviant and standard responses. The observed cluster weights were then compared to this permutation distribution. A cluster was considered statistically significant if its weight exceeded 95 % of the values in the permutation distribution, corresponding to a p-value of 0.05. The p-values of all observed clusters were corrected for multiple comparisons using the Benjamini-Hochberg false discovery rate (FDR) method (Benjamini and Hochberg, 1995).

Given that clusters of significant differences between deviant and standard ABRs were only found for the low-frequency chirp, all subsequent analyses focused solely on responses to the low-frequency chirp. To determine whether the significant differences between the waveforms of deviant and standard ABRs were due to variations in ABR peak-to-peak amplitude and peak V latency, additional statistical tests were conducted. These parameters were calculated within predefined analysis windows. The peak-to-peak amplitude was measured within a window comprising all significant clusters, as identified by the cluster- based permutation analysis (2.7-9.2 ms post-stimulus peak). For peak V latency, a later window onset was selected (5-9.2 ms post-stimulus peak) to focus exclusively on wave V and minimise data variability. Both peak-to-peak amplitude and peak V latency were confirmed to be normally distributed using Lilliefors tests.

To assess the effects of stimulus probability (deviant, standard) and stimulus repetition rate (40 Hz, 33 Hz, 20 Hz) on ABR peak-to-peak amplitude and peak V latency, a two-way analysis of variance (ANOVA) was performed, considering potential interactions between the two factors. In case of significant ANOVA results, post hoc t-tests were carried out, with adjustments for multiple comparisons using the Benjamini-Hochberg FDR method. Additionally, Cohen’s *d* was calculated for each significant comparison to provide a measure of effect size (Cohen, 1988).

The second part of this study explored how ABR amplitude and peak latency are modulated by deviance detection as the repetition number of the standard stimulus increases following a deviant (Fig. 3a). Sequences containing a deviant and at least three consecutive standard responses were extracted from the oddball recording, and the deviant as well as the first, second and third standard response were averaged per participant (with trial numbers ranging between 683 and 723). The effects of stimulus role (deviant, standard 1, standard 2, standard 3) and repetition rate (40 Hz, 33 Hz, 20 Hz) on ABR peak-to-peak amplitude and peak V latency were assessed using a two-way ANOVA, accounting for potential interactions. When significant results were identified in the ANOVA, post-hoc t-test comparisons were performed with Benjamini-Hochberg FDR correction, and Cohen’s *d* was calculated for significant t-test outcomes to quantify the effect size. The data and code used for this study are freely available on G-Node (DOI: 10.12751/g-node.bh5dfs).

## 3. Results

### Deviance detection modulates ABR amplitude and latency

To investigate the effects of deviance detection in the brainstem, ABRs were recorded using a low- and a high-frequency chirp stimulus, presented in an oddball paradigm (Fig. 1b). To explore the effect of stimulus repetition rate, three different rates were tested: 40 Hz, which is a commonly used rate in ABR research, and two slower rates of 33 Hz and 20 Hz. A cluster-based permutation analysis was performed to determine which ABR components differed significantly between deviant and standard responses. For the low-frequency chirp, a significant cluster was identified between 8 and 10 ms after basilar membrane activation across all stimulus repetition rates (40 Hz: p = 0.029; 33 Hz: p = 0.020; 20 Hz: p = 0.031), coinciding with the falling slope of ABR wave V (Fig. 2a). Additionally, at the fastest repetition rate of 40 Hz, a second, earlier significant cluster was observed between 2.7 and 4.1 ms (p < 0.001) (Fig. 2a, top). In contrast, no significant clusters were detected for any of the high-frequency chirp responses (Fig. 2b).

**Figure 2:**
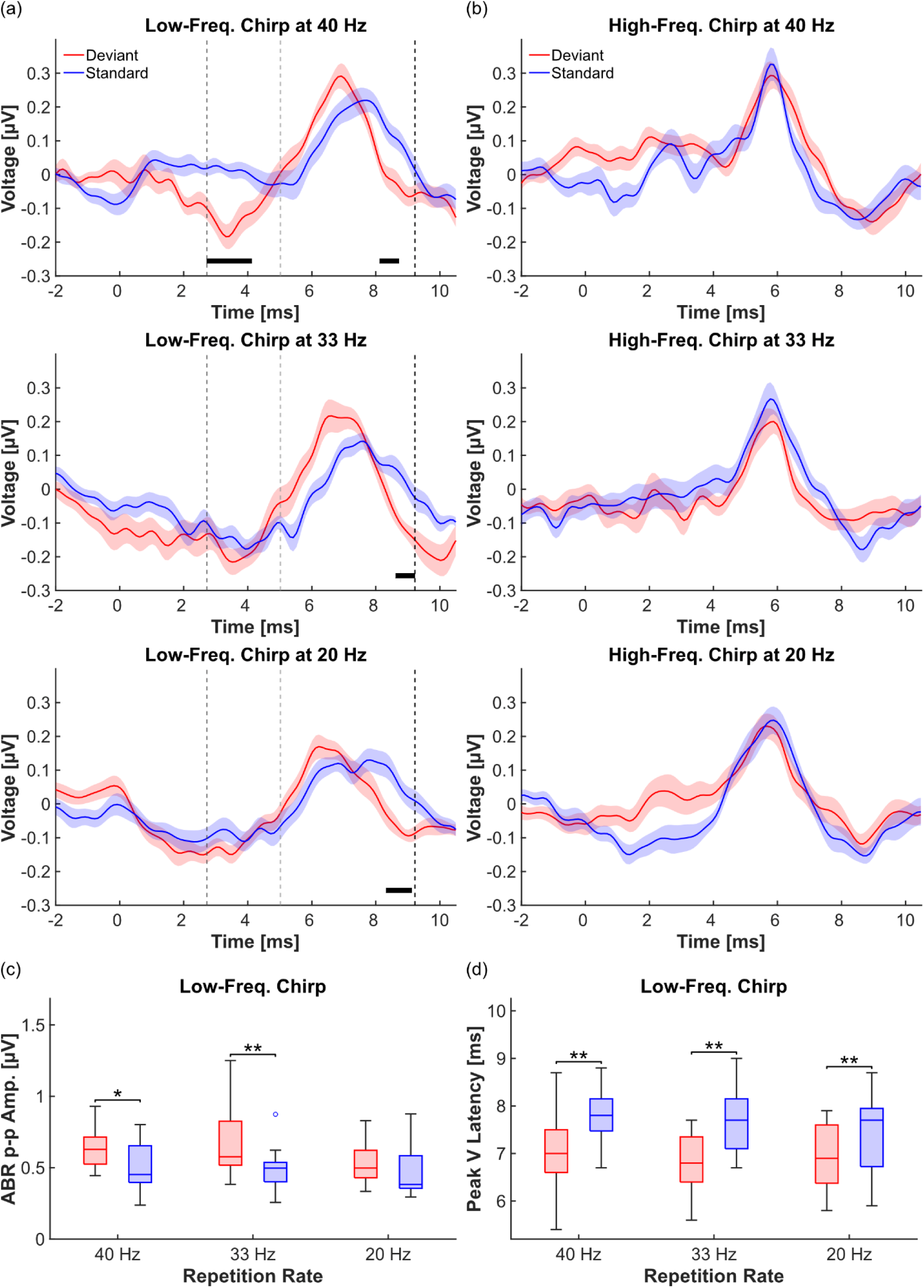
Deviance detection modulates ABR amplitude and latency (n = 17 subjects). (a) Grand average ABRs elicited by a low-frequency chirp, presented as deviant (red) and standard (blue) at three different stimulus presentation rates (40 Hz, top; 33 Hz, middle; 20 Hz, bottom). The black bars below the ABRs indicate significant differences between deviant and standard waveforms, identified through a cluster-based permutation analysis. Vertical dashed lines mark the time windows used for subsequent amplitude and latency analyses: from the dark grey (first) line to the black (third) line for amplitude analysis and from the light grey (second) line to the black (third) line for peak latency analysis. Shaded areas represent the standard error of the mean. (b) As in (a), but the stimulus was a high-frequency chirp. The cluster-based permutation analysis revealed no significant differences between deviant and standard responses. (c) Boxplots representing the peak-to-peak values of low-frequency chirp ABRs, where red boxes represent deviant responses, and blue boxes represent standard responses. Peak-to-peak values are compared within each stimulus repetition rate. (d) As in (c), but boxplots represent ABR peak V latency.

To identify the response parameters responsible for the significant clusters observed in the low-frequency responses, ABR peak-to-peak amplitude and ABR peak V latency were calculated and compared using two-way ANOVAs, with stimulus probability (deviant, standard) and stimulus repetition rate (40 Hz, 33 Hz, 20 Hz) as factors. The dependent variables were calculated within predefined time windows, indicated by the vertical dashed lines (Fig. 2a,b). The window for peak-to-peak calculation covered all significant clusters (from dark grey line, 2.7 ms, to black line, 9.2 ms) and the latency window was set to only cover peak V (from light grey line, 5.0 ms to black line, 9.2 ms). For the low-frequency chirp, the ANOVA revealed a significant effect of stimulus probability on both ABR peak-to-peak amplitude (p = 4.9*10⁻⁴) and peak V latency (p = 2.1*10⁻⁶). Post hoc t-tests comparing deviant and standard responses demonstrated that peak-to-peak amplitude was significantly different at the two faster repetition rates of 40 Hz (p = 0.010, Cohen’s d = 0.84) and 33 Hz (p = 0.006, Cohen’s d = 0.94), with large effect sizes (Fig. 2c). Peak V latency was significantly different across all three tested repetition rates (40 Hz: p = 0.005, Cohen’s d = 1.11; 33 Hz: p = 0.007, Cohen’s d = 1.27; 20 Hz: p = 0.007, Cohen’s d = 0.62), with large effect sizes for the two faster rates and medium effect size for the slower rate (Fig. 2d).

### ABR deviance detection is rapid and differentially affects response parameters

SSA studies in animals have demonstrated that deviance detection emerges rapidly in subcortical areas (Malmierca et al., 2009; Antunes et al., 2010; Bäuerle et al., 2011; Ayala and Malmierca, 2012). To investigate whether the same phenomenon can be observed in humans, we reanalysed the low-frequency chirp data obtained in the oddball paradigm to examine how the difference between deviant and standard response develops as the index of standard repetitions increases (Fig. 3a).

**Figure 3:**
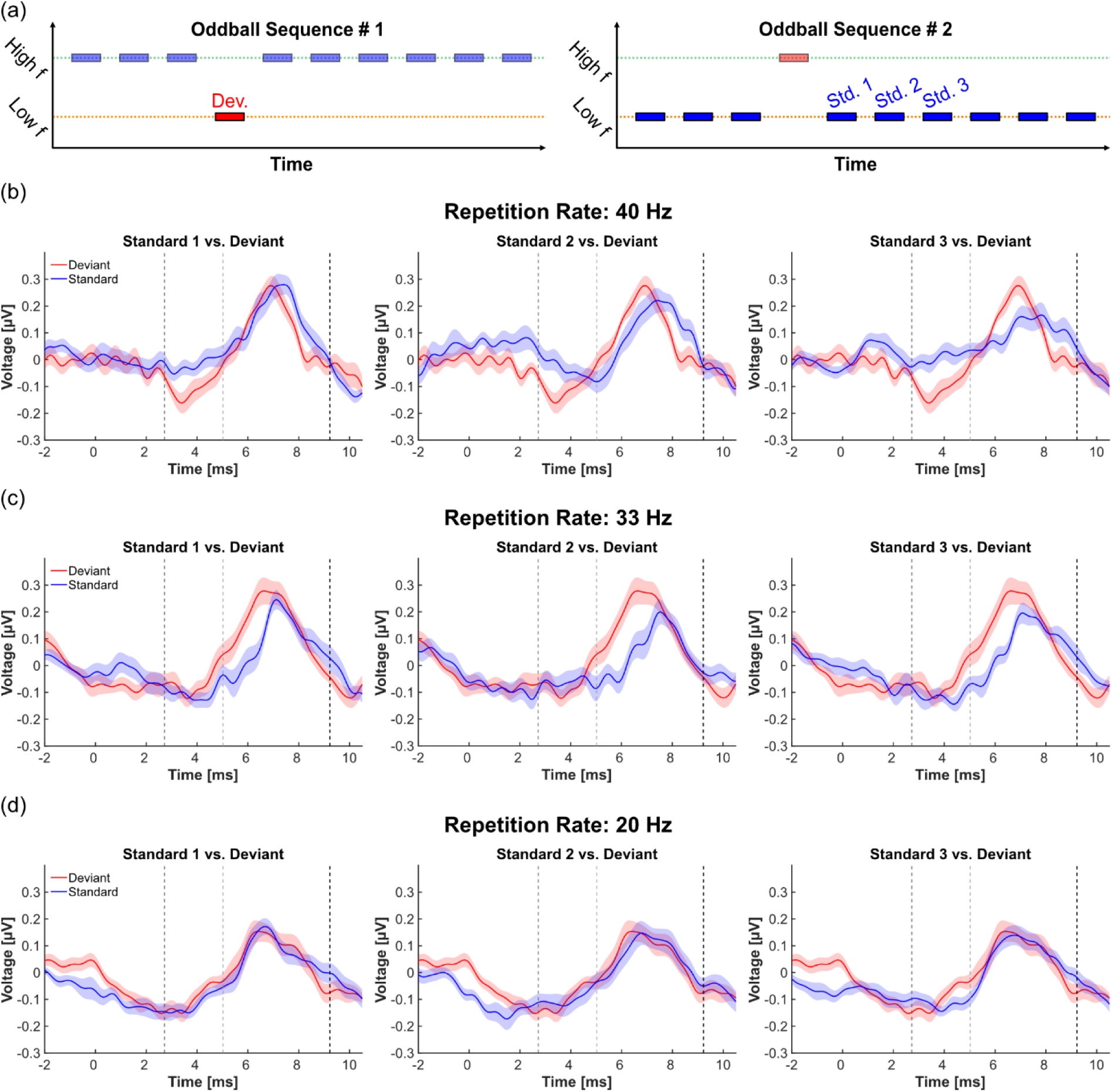
ABR waveform changes with increasing standard repetition (n = 17 subjects). (a) Schematic representation of the responses illustrated in this figure. Focusing on the low- frequency chirp ABRs, the first three standard responses following a deviant (Std. 1 - Std. 3, blue) are individually compared to the deviant ABRs of the same low-frequency chirp stimulus (Dev., red). (b) Grand average ABRs elicited by a low-frequency chirp. Comparison between the deviant ABR (red) and the responses to standards 1-3 (blue), presented at a stimulus repetition rate of 40 Hz. Vertical dashed lines indicate the time windows used for subsequent amplitude and latency analyses: from the dark grey (first) line to the black (third) line for amplitude analysis, and from the light grey (second) line to the black (third) line for peak latency analysis (Fig. 4). Shaded areas represent the standard error of the mean. (c) As in (b), but with a stimulus repetition rate of 33 Hz. (d) As in (b), but with a stimulus repetition rate of 20 Hz.

Specifically, we compared the deviant response to the first three standard responses following a deviant. ABR peak-to-peak amplitude and peak V latency were calculated within the same time windows as in the first section of the study (Fig. 3b-d). A two-way ANOVA with the factors of stimulus role (deviant, standard 1, standard 2, standard 3) and repetition rate (40 Hz, 33 Hz, 20 Hz) revealed a significant effect of stimulus role on both ABR peak-to-peak amplitude (p = 0.036) and peak V latency (p = 5.7*10⁻⁴). Subsequent post-hoc t-tests compared the deviant response with each of the standard responses within the same repetition rate. For the 40 Hz rate, a significant difference with a large effect size was found between the deviant and the third standard repetition (p = 0.026, Cohen’s d = 0.92), indicating a significant repetition suppression effect after no more than three standard repetitions (Fig. 4a, left). Conversely, while a visual inspection suggests a decline in response amplitude at the 33 Hz rate (Fig. 3c), the effect was not statistically significant (Fig. 4a, middle). Finally, no significant effect was observed for the 20 Hz repetition rate (Fig. 3d; Fig. 4a, right).

**Figure 4:**
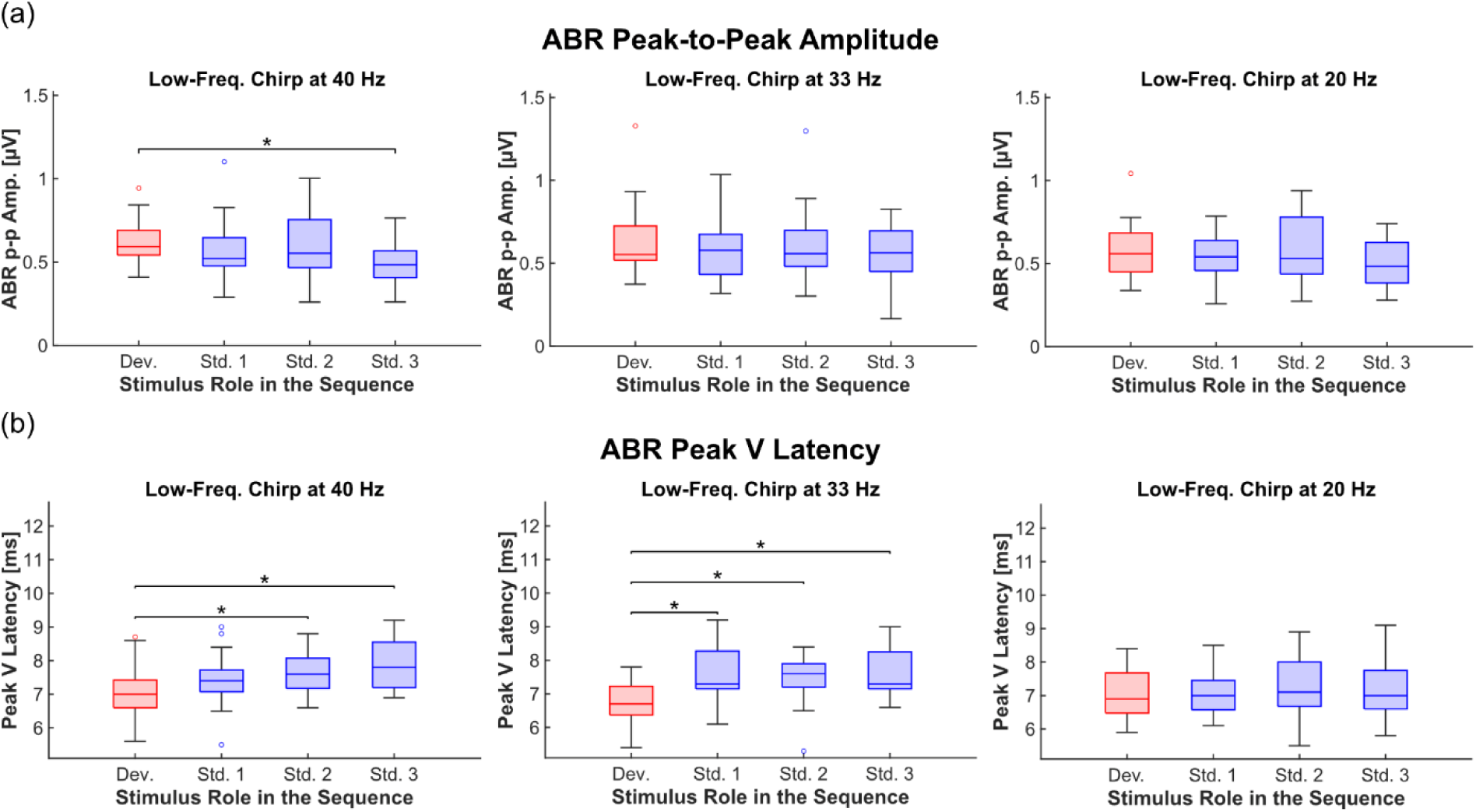
ABR deviance detection is rapid and differentially affects response parameters (n = 17 subjects). (a) Boxplots showing the peak-to-peak amplitude of ABRs in response to the low-frequency chirp. The peak-to-peak values for the deviant response (red) are compared to those of the first three standard responses following a deviant (blue). The experiment was conducted at three different stimulus repetition rates: 40 Hz (left), 33 Hz (middle), and 20 Hz (right). The data corresponds to the responses illustrated in Fig. 3. (b) As in (a), but the boxplots represent ABR peak V latency.

Post hoc analysis of ABR peak V latency revealed significant differences between the deviant response and the second (Dev. vs. Std. 2: p = 0.026, Cohen’s d = 0.78; Dev. vs. Std. 3: p = 0.020, Cohen’s d = 0.99) and even the first (Dev. vs. Std. 1: p = 0.021, Cohen’s d = 1.23; Dev. vs. Std. 2: p = 0.025, Cohen’s d = 0.95; Dev. vs. Std. 3: p = 0.036, Cohen’s d = 1.27) standard repetition for the 40 Hz and 33 Hz stimulation rates, respectively (Fig. 4b, left and middle).

For the 20 Hz stimulus presentation rate, the post hoc analysis did not reveal significant differences between the deviant and any of the standard responses (Fig 4b, right), indicating that the latency shift observed at this stimulus presentation rate in the first part of the study (Fig. 2d) requires more than three standard repetitions to emerge.

## 4. Discussion

This study employed ABRs to examine subcortical auditory deviance detection in humans. Low-frequency chirps presented as deviants evoked significantly larger and faster ABRs compared to when the same stimuli were used as standards. These effects were detectable within less than 10 ms after basilar membrane activation. A comparison of the deviant response with the three standard responses immediately following a deviant revealed that ABR differences emerged within up to three standard repetitions at the fastest stimulus presentation rate. The data further demonstrate that deviance detection differentially influences response amplitude and latency. Additionally, the findings indicate that subcortical deviance detection is stimulus-specific – as no significant differences were observed between deviant and standard responses for high-frequency chirps – and becomes more pronounced at higher stimulus presentation rates.

These findings are consistent with animal research that has investigated SSA at the single-unit level in the IC of rats, where deviant stimuli were shown to elicit not only more frequent but also earlier single-neuron responses compared to standard stimuli (Malmierca et al., 2009; Zhao et al., 2011). Moreover, studies in rodents have demonstrated that subcortical SSA occurs within less than 10 standard repetitions (Malmierca et al., 2009; Antunes et al., 2010; Bäuerle et al., 2011; Ayala and Malmierca, 2012) and is dependent on the stimulus presentation rate (Malmierca et al., 2009; Antunes et al., 2010; Zhao et al., 2011; Patel et al., 2012).

The current study bridges a longstanding gap between animal and human research concerning the latency of early deviance detection. While animal studies have shown SSA in the IC with latencies of less than 15 ms (Pérez-González et al., 2005; Malmierca et al., 2009), the earliest deviance detection in humans was so far measured at latencies around 25 to 30 ms after stimulus onset (Slabu et al., 2010; Althen et al., 2011; Grimm and Escera, 2012; Shiga et al., 2015). Our ABR data now show that deviance detection with latencies below 10 ms also occurs in humans, suggesting similar temporal dynamics in subcortical deviance detection processing across species. Interestingly, our findings also indicate that significant differences between deviant and standard ABRs appear earlier in the signal when the stimulus presentation rate is high. Deviant and standard ABRs consistently differed between approximately 8 ms and 9.2 ms after basilar membrane activation across all tested repetition rates. However, at the highest rate (40 Hz), an additional cluster of significant waveform differences was observed between 2.7 and 4.1 ms (Fig. 2). This earlier cluster coincides temporally with neural activity in the cochlear nucleus and superior olivary complex (Buchwald and Huang, 1975; Henry, 1979; Parkkonen et al., 2009), suggesting that at high repetition rates, deviance detection may modulate the responsiveness of brainstem nuclei even below the IC. These findings align with previous studies on deviance detection in bat ABRs (Wetekam et al., 2022, 2024). However, unlike the ABRs reported in these animal studies, the human ABRs presented here lack prominent waves I–IV, which are typically associated with the activity of auditory nuclei below the IC (Buchwald and Huang, 1975; Henry, 1979; Parkkonen et al., 2009). Instead, the data are dominated by ABR wave V, mainly generated in the IC (Buchwald and Huang, 1975; Henry, 1979; Caird and Klinke, 1987; Parkkonen et al., 2009; Land et al., 2016). Consequently, the influence of deviance detection on subcollicular nuclei should be interpreted cautiously. Future studies employing stimuli that elicit larger ABR waves I–IV in humans would be valuable to further explore this possibility.

The finding that significant deviance detection emerges after only a few standard repetitions is consistent with previous research in animals (Malmierca et al., 2009; Antunes et al., 2010; Bäuerle et al., 2011; Ayala and Malmierca, 2012). However, to our knowledge, this study presents the first evidence of a differential influence of deviance detection on subcortical response amplitude and latency. While amplitude differences between deviant and standard responses only become evident after three or more standard repetitions, latency differences can be observed as early as the first standard repetition at certain stimulus presentation rates. One possible explanation for this phenomenon is that the shorter latency observed for deviant responses, compared to standard responses, may not be solely attributable to an increasing delay of the standard response with each repetition (repetition delay). Instead, it could involve an additional reduction in response latency when a deviant is detected. Testing this hypothesis in future studies would be valuable and could be explored further using control experiments like the many-standards sequence (Schröger and Wolff, 1996).

Another open question that arises is why significant deviance detection was only elicited by the low-frequency chirp. Similar findings have been observed in bat ABRs (Wetekam et al., 2024) and human FFRs (Slabu et al., 2012), but the underlying reasons remain unclear. In our study, one possible explanation for this discrepancy may stem from research on repetition suppression. These studies have shown that repetition suppression, a key mechanism in deviance detection, is more commonly observed for familiar sounds compared to unfamiliar ones (Henson et al., 2000; Müller et al., 2013; Segaert et al., 2013). The spectrum of our low-frequency chirp (500-2000 Hz) comprises most of the frequencies typically encountered in daily life, making it a potentially more familiar stimulus compared to the high-frequency chirp (2000-8000 Hz). Furthermore, high-pitched sounds are often associated with alarm signals (Zeskind and Collins, 1987; Haas and Edworthy, 1996; Schwartz and Gouzoules, 2019), which may gain behavioural relevance when perceived repeatedly. Consequently, the primary goal of processing high-frequency stimuli might not be to maximise efficiency, but rather to maintain an effective alarm signal that does not diminish with repeated exposure. These hypotheses should be addressed in future research, together with a more detailed investigation of the specific stimulus parameters required to elicit mismatch responses in ABRs, e.g. identifying the frequency threshold below which deviance detection becomes measurable.

Beyond stimulus frequency, other parameters of stimulation, such as the stimulus repetition rate, have also been shown to significantly influence subcortical deviance detection in animals (Malmierca et al., 2009; Antunes et al., 2010; Zhao et al., 2011; Patel et al., 2012). Our data corroborate this dependency by suggesting that mismatch responses in ABRs decrease with the stimulus presentation rate. This finding makes it unlikely that significant effects would have been observed at repetition rates below 10 Hz, potentially explaining why two earlier studies failed to detect an effect of stimulus probability on ABRs (Slabu et al., 2010; Althen et al., 2011). In those studies, the authors used lower stimulus repetition rates of approximately 10 Hz (Slabu et al., 2010) and 3 Hz (Althen et al., 2011), along with higher deviant probabilities of 1/5 (Slabu et al., 2010) and 1/7 (Althen et al., 2011). Lastly, it is crucial to emphasis the careful consideration of analysis parameters when investigating deviance detection in ABRs. At the high repetition rates required to elicit significant effects, artefacts from overlapping responses can confound results. Specifically, when the ABR overlaps with the MLR elicited by the preceding stimulus, changes in amplitude may not exclusively reflect stimulus probability but may instead represent differences in MLR components evoked by earlier stimuli. Therefore, it is essential to apply filter and baseline correction settings that ensure the observed effects are independent of responses to preceding stimuli and thus solely represent deviance detection in the brainstem.

## Conclusion

This study revealed that subcortical auditory deviance detection in humans is both rapid and stimulus-specific, with significant modulation of ABR amplitude and latency occurring within just a few stimulus repetitions. The identification of mismatch responses at latencies below 10 ms helps to bridge the gap between human and animal research, indicating similar temporal dynamics across species. Moreover, the study highlights the value of ABR recordings for investigating subcortical auditory deviance detection, due to its excellent temporal resolution in comparison to other non-invasive methods.

## Acknowledgements

This work was funded by the Deutsche Forschungsgemeinschaft [KO 987/14-1].

## Conflict of Interest

The authors declare no competing interests.

## Author contributions

Study design: Johannes Wetekam, Nell Gotta, Luciana López-Jury, Julio Hechavarría, Manfred Kössl

Experiments: Johannes Wetekam, Nell Gotta

Analysis, and original draft of the manuscript: Johannes Wetekam

Review and editing of the original draft: Johannes Wetekam, Nell Gotta, Luciana López-Jury, Julio Hechavarría, Manfred Kössl

## Abbreviations

ABR: Auditory brainstem response
ANOVA: Analysis of variance
FDR: False discovery rate
FFR: Frequency-following response
fMRI: Functional magnetic resonance imaging
IC: Inferior colliculus
MLR: Middle-latency response
SSA: Stimulus-specific adaptation

## Notes

### Competing Interest Statement

The authors have declared no competing interest.

### Summary of Updates

The article has been updated based on peer-review feedback. Notably, the introduction has been thoroughly revised for clarity and coherence. Data discussion previously included in the results section has been removed to streamline the presentation, and the discussion section has been refined to enhance focus and overall clarity. The presented data remain unchanged.

